# Predicting the Trend of SARS-CoV-2 Mutation Frequencies Using Historical Data

**DOI:** 10.1101/2023.12.19.572480

**Authors:** Xinyu Zhou, Kevin Hu, Minmin Pan, Yajie Li, Chi Zhang, Sha Cao

**Affiliations:** Center for Computational Biology and Bioinformatics, Department of Medical and Molecular Genetics, School of Medicine, Indiana University, Indianapolis, IN; Department of Computer Science, Indiana University Bloomington, Bloomington, IN; Carmel High School, Carmel, IN; Department of Biostatistics and Health Data Science, School of Medicine, Indiana University, Indianapolis, IN

## Abstract

As the SARS-CoV-2 virus rapidly evolves, predicting the trajectory of viral variations has become a critical yet complex task. A deep understanding of future mutation patterns, in particular the mutations that will prevail in the near future, is vital in steering diagnostics, therapeutics, and vaccine strategies in the coming months.

In this study, we developed a model to forecast future SARS-CoV-2 mutation surges in real-time, using historical mutation frequency data from the USA. To improve upon the accuracy of traditional time-series models, we transformed the prediction problem into a supervised learning framework using a sliding window approach. This involved breaking the time series of mutation frequencies into very short segments. Considering the time-dependent nature of the data, we focused on modeling the first-order derivative of the mutation frequency. We predicted the final derivative in each segment based on the preceding derivatives, employing various machine learning methods, including random forest, XGBoost, support vector machine, and neural network models, in this supervised learning setting. Empowered by the novel transformation strategy and the high capacity of machine learning models, we witnessed low prediction error that is confined within 0.1% and 1% when making predictions for future 30 and 80 days respectively. In addition, the method also led to a notable increase in prediction accuracy compared to traditional time-series models, as evidenced by lower MAE, and MSE for predictions made within different time horizons. To further assess the method’s effectiveness and robustness in predicting mutation patterns for unforeseen mutations, we categorized all mutations into three major patterns. The model demonstrated its robustness by accurately predicting unseen mutation patterns when training on data from two pattern categories while testing on the third pattern category, showcasing its potential in forecasting a variety of mutation trajectories.

To enhance accessibility and utility, we built our methodology into an R-shiny app (https://swdatpredicts.shinyapps.io/rshiny_predict/), a tool with potential applicability in studying other infectious diseases, thus extending its relevance beyond the current pandemic.

## 1 Introduction

The COVID-19 pandemic, caused by the severe acute respiratory syndrome coronavirus 2 (SARS-CoV-2), continues to pose significant global challenges. As of September 2023, over 770 million people have been infected, resulting in the tragic loss of around seven million lives [1]. A pivotal aspect of the pandemic has been the emergence of numerous SARS-CoV-2 variants with different properties such as increased transmissibility, changes in disease severity, and varied responsiveness to vaccines and treatments [2–5], significantly affecting the effectiveness of public health interventions [3, 5–7]. Hence, understanding the mutation dynamics of the virus is critical in navigating the ongoing crisis efficiently [8]. Mutations in the viral genome occur due to errors in RNA replication processes, resulting in variations such as base substitutions, insertions, and deletions [9, 10]. With ongoing transmission, these microscopic changes can be replicated in subsequent rounds of infection, and potentially alter the virus’s characteristics, further influencing the efficacy of diagnostic tools, vaccines, and therapies [10].

To better understand the virus mutation dynamics, the global scientific community has collected a substantial amount of genomic data, with about 16 million SARS-CoV-2 sequences currently available through the Global Initiative on Sharing All Influenza Data (GISAID) [11]. This extensive database has catalyzed a new age in viral genomic research, fostering near realtime surveillance of the pandemic and profoundly influencing public health policies [11, 12]. For instance, researchers have developed real-time monitoring systems to track the emergence and spread of new variants globally [1, 13, 14]. Researchers have also relied on the mutation fingerprints for studying real-time molecular epidemiology [15–18]. Several recent reports describe new methods on utilizing machine learning and computational biology to predict potential future mutations and their implications, and forecast the potential impact of mutations on the trajectory of the pandemic [19, 20]. The differentiation of the SARS-CoV-2 virus according to geographic locations [21] and racial differences, as well as the impacts it has on humans post-infection [22], also constitute hotspots in research.

Despite the breadth of ongoing scientific investigation, a critical challenge remains to be the accurate foresight of dominant variants in the imminent future. Presently, a myriad of forecasts exists concerning SARS-CoV-2 variants [23–26]. Although insightful, these methods may fall short of predicting variants that emerge in the future because of the complexity of host–virus interaction and the plasticity of virus biology [27]. In particular, the current methods primarily focus on predicting macro-level variants, potentially overlooking the subtleties of individual mutations that significantly impact the virus’s behavior. Additionally, these methods do not provide real-time application which is crucial for public health responses.

While variants have gained significant attention, it is their constituent mutations that directly influence the effectiveness of diagnostic tools and vaccines. This raises a compelling proposition: rather than focusing solely on predicting dominant variants, we may shift our attention to the underlying mutations. By extrapolating the future frequencies of individual mutations and discerning dominant trends, we gain deeper insights into the genomic dynamics of the virus in real-time, which is an approach previously unexplored.

In this study, we focus on predicting the virus genomic dynamics by focusing on individual mutations. We propose to utilize historical mutation frequency data from US patients in GISAID to forecast future mutation frequency patterns in real-time, while also determining the maximum time horizon within which mutation frequencies can be accurately predicted with minimal deviation. Recognizing the limitations of traditional time series-based models, we have devised a data transformation strategy, thereby converting a time series modeling challenge into a supervised learning task. This innovation enables the implementation of advanced machine learning methods to harness their critical predictive capacities. Our research offers several key contributions: 1) Studying the virus genomic dynamics at the unit of individual mutations, an approach not previously pursued. 2) A novel data transformation method designed to harness machine learning capabilities for mutation frequency predictions. 3) Modeling based on the first-order derivative of mutation frequency, enhancing prediction accuracy and capturing temporal relationships. 4) The establishment of evaluation metrics for various methods, enabling a comprehensive assessment of their effectiveness.

Throughout this article, we employ the term “mutation” to denote base-level changes comparing to the Wuhan-Hu-1 reference sequence (GenBank accession: NC 045512.2).

## 2 Results

### 2.1 Problem formulation

Assume there are in total *K* genomic locations whose daily mutation rates have been observed from day 1 to *N* in a certain population. For a specific mutation *k*, we denote the mutation frequency for day *i* as *f*_*ki*_, where *i* = 1, …, *N*; *k* = 1, …, *K*. Our objective is to learn a forecasting function *g* that can accurately predict the mutation rate *f*_*k,N*+1_, …, *f*_*k,N*+*T*_ for a specific mutation *k* on future *T* days, based on the historical rates observed from day 1 to day *N*. Formally, we define the function as:

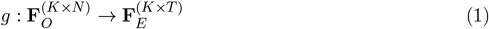

This function takes as input the matrix of observed mutation rates **F**_*O*_ *∈ ℛ*^*K×N*^, where each element represents the rate of mutation *k* on day *i*, and outputs a matrix **F**_*E*_ *∈ ℛ*^*K×T*^, with each row corresponding to the predicted rate of mutation for each of the *K* locations for the future *T* days. The challenge is to design and train *g* such that it minimizes a loss function 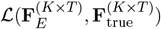, where 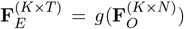, and 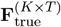 is the underlying truth. This loss function quantifies the accuracy of the predictions, and the goal is to find the right *g* that could minimize this loss over the training dataset. Note that depending on the choice of the model, *g* may vary across different *k* such as time series model. This mathematical formulation sets a clear objective for the learning task, focusing on predicting future mutation rates based on historical data. Clearly, there are three key aspects with this formulation: constructing the training and testing dataset; choice of family of prediction function *g*, and the evaluation metrics.

### 2.2 Harnessing the power of machine learning methods in modeling mutation sequence data

In section 2.1, our task involves forecasting based on time. While traditional time-series models and curve fitting techniques like spline methods are common choices, they may struggle with the complex, hard-to-learn patterns of mutation frequency. As we’ll demonstrate, these methods don’t always yield the best results in our context. Nowadays, machine learning has become increasingly effective in deciphering complex biological patterns. Leveraging this, we introduce an innovative approach: transforming each full-length temporal mutation frequency sequence into shorter, consecutive segments. This method, known as sliding window dissection (**SWD**), redefines a time series problem as a supervised prediction challenge. With SWD, each segment’s last data point becomes the response variable, while the preceding points serve as predictors. This setup integrates temporal dependencies into the supervised learning framework. It allows for more sophisticated modeling of nonlinear relationships and interactions using advanced machine learning techniques. Models like random forest, neural networks, support vector machines (SVM), and XGBoost are adept at learning intricate patterns and adapting to evolving dynamics. Consequently, they provide reliable predictions even amid nonlinear and irregular trends in the data. Most importantly, a key advantage of this transformation strategy is its ability to integrate mutation frequency data from different genomic locations into a single predictive model. This holistic approach could enhance the accuracy and robustness of our predictions, offering a more comprehensive understanding of the virus’s evolution.

Training the aforementioned advanced models is a pivotal step, highly dependent on the training dataset (Figure 1A). From GISAID, we collected the mutation rate data spanning a total of 1,130 days, for a total of 87 genomic locations. Details on the 87 genomic locations is illustrated in Methods and Materials section. Figure 1B illustrates how we constructed our training data using a sliding window approach. In this method, for a segment of length *d* + 1 within the range [1, …, *N*], specifically [*i*, …, *i* + *d*], we treat *f*_*k,i*_, *f*_*k,i*+1_, …, *f*_*k,i*+*d-*1_ as predictors (denoted as **x**_*ki*_) and *f*_*k,i*+*d*_ as the response (denoted as *y*_*ki*_), where *d* is the window size. By iterating over all values of *k* and *i*, we obtain all possible sliding segments (**x**_*ki*_, *y*_*ki*_), *k* = 1, …, *K, i* = 1, …, *N - j - d*. A portion of these segments forms our training data, helping us identify the most effective prediction model. The remaining segments are used as testing data to evaluate the methods’ performance. This method prepares our data for application in various machine learning models. Each model can be trained using the training data and assessed using the testing data. It’s important to note that, due to the temporal nature of the data, all training data segments must precede the testing data segments in time. This ensures the validity and reliability of our forecasting approach.

**Figure 1:**
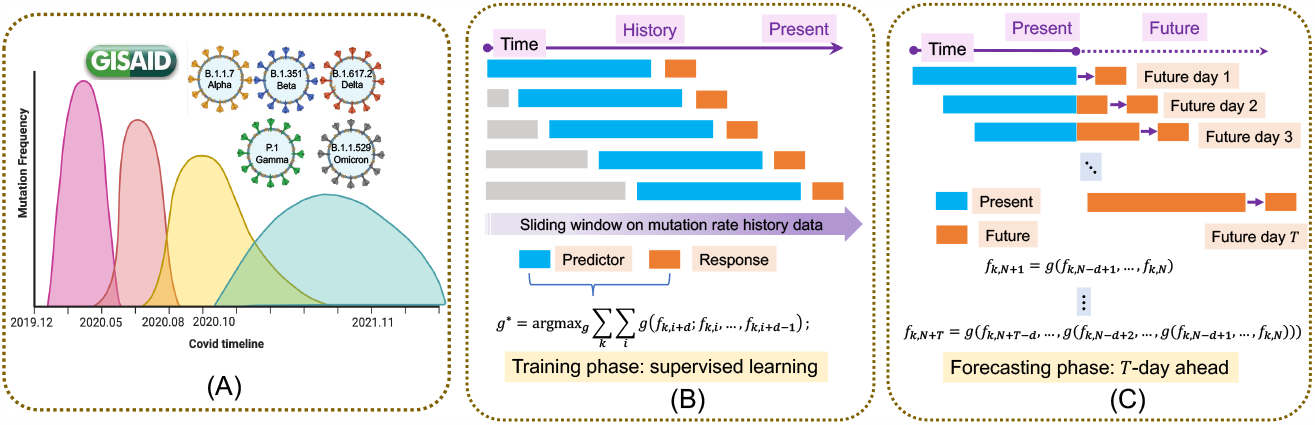
Schematic outline of our method. A) Different COVID variants have appeard so far, each with different combinations of mutation types with varied mutation frequencies observed in the population. To forecast the mutation rate changes, our prediction model consists of a training phase (B) and forecasting phase (C). B) In the training phase, for all mutation, their historical mutation rate is calculated, and using a sliding window approach, we could apply machine learning model to predict the last mutation pattern using preceding ones. C) In the forecasting phase, future mutation rates is predicted day by day with a recursive fashion.

Figure 1C illustrates the forecasting process for the mutation rate of a specific location over the next *T* days. The prediction begins with 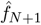, using observed rates {*f*_*N-d*+1_, *…, f*_*N*_}. The process then recursively predicts 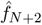 using 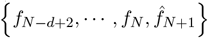 as inputs, continuing in this manner until 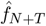 is forecasted. A key advantage of our approach is the ability to combine mutation patterns from various types to create a unified prediction model. In contrast, while traditional time series models excel at capturing periodic or seasonal trends by training on individual sequences, they may fall short in integrating mutation frequency data from diverse types into a cohesive prediction model. This limitation highlights the flexibility and comprehensiveness of our forecasting method.

In our data engineering process, we initially used a sliding window method to create **x**_*ki*_. However, this method overlooked the sequence order in **x**_*ki*_ = (*f*_*ki*_, *f*_*k,i*+1_, …, *f*_*k,i*+*d-*1_). To address this, we shifted our focus from modeling daily mutation rates to their first-order derivatives. Consequently, our new predictor and response variables are 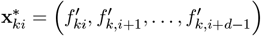 and 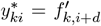, where 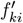 represents the first-order derivative of the mutation rate at day *i* for location *k*. Details can be found in the Methods and Materials section. This approach to first-order derivatives allows us to more effectively capture the dependencies within the predictor variables. Another significant benefit is the reduction of auto-correlation among predictors. This auto-correlation, arising from overlapping data points in consecutive sequences, can skew the learning process of the model and lead to biased, overly optimistic evaluations. By using first-order derivatives and introducing lag variables, we decrease the dependence between consecutive observations, enhancing the robustness and reliability of our modeling.

In our study, we explored a wide range of prediction techniques on transformed data. These include SVM with linear (S_*L*_) and radial (S_*R*_) kernels, random forest (RF), XGBoost (XG), forward neural network (FNN), and linear regression (LR) as a straightforward, interpretable benchmark. This selection spans various modeling strategies, blending linear and non-linear, parametric and non-parametric models, alongside ensemble methods and neural networks. This methodological diversity ensures a comprehensive analysis, as the strengths and limitations of each method are balanced by others, boosting the credibility and broad applicability of our results. We refer to these as supervised learning methods, denoted by SWD+X, where X represents one of S_*L*_, S_*R*_, RF, XG, FNN, LR. In contrast, we also evaluated traditional methods that have been used to model time series data, such as ARIMA (ARI), Prophet (PRO), *k*-nearest neighbour (*k*NN), cubic-spline (C_*S*_), and B-spline (B_*S*_). ARIMA and Prophet are autoregressive models typically used in time series analysis, while *k*NN and spline methods are more functional, designed to fit curves to time series data. For convinience, these were all referred to as time series learning methods. A key part of our analysis was comparing these traditional models against the supervised learning methods.

### 2.3 Performance evaluations of supervised learning and time series learning methods

This section first evaluates supervised and time series learning approaches using Mean Square Error (MSE) and Mean Absolute Error (MAE). These metrics assess the deviation between predicted and actual mutation frequencies in test data. For supervised learning, we split the whole time frame at a specific date, *t*_0_. Here, we set *t*_0_ = 500. Data before *t*_0_ served as training material, covering all genomic locations. We then tested predictions for days 1,7,14,30,80 beyond *t*_0_, comparing them against the actual mutation rates. Time series methods also used *t*_0_ as a dividing point. Here, each mutation rate sequence was learned independently, using data before *t*_0_ for training. We then predicted mutation rates after *t*_0_ and contrasted them with the actual values. For more information, refer to the Methods and Materials section. Importantly, *t*_0_ was not set close to the end of the time series. This choice avoids periods where the mutation rate curve is mostly flat and near zero, as seen in Supplementary Figure 1, ensuring a more robust evaluation of the methods under varied mutation frequencies.

Figure 2 clearly shows that supervised learning methods (indicated by cold color bubbles) outperform time series approaches (warm color bubbles), achieving lower mean absolute error (MAE) and mean squared error (MSE). Within supervised learning, random forest and SVM (linear/radial) emerge as the most accurate, recording the lowest MAE and MSE on average. These methods consistently surpass all time series methods across all time frames. Notably, linear regression, under our data transformation framework, matches these high standards, outperforming time series methods in predicting future days 1, 7, 14, and 80, and sometimes even excelling XGBoost and FNN. In the time series category, none can match supervised learning methods for shorter predictions (future days 1, 7, 14). However, for longer forecasts (future days 30, 80), ARIMA, the best among all selected time series learning methods, shows comparable or slightly superior performance to linear regression, XGBoost, and FNN, but its performance is still much worse than random forest and SVM. It’s evident that with increasing prediction time frames, supervised learning methods face escalating errors, a result of error accumulation during the recursive prediction phase (see Figure 1).

**Figure 2:**
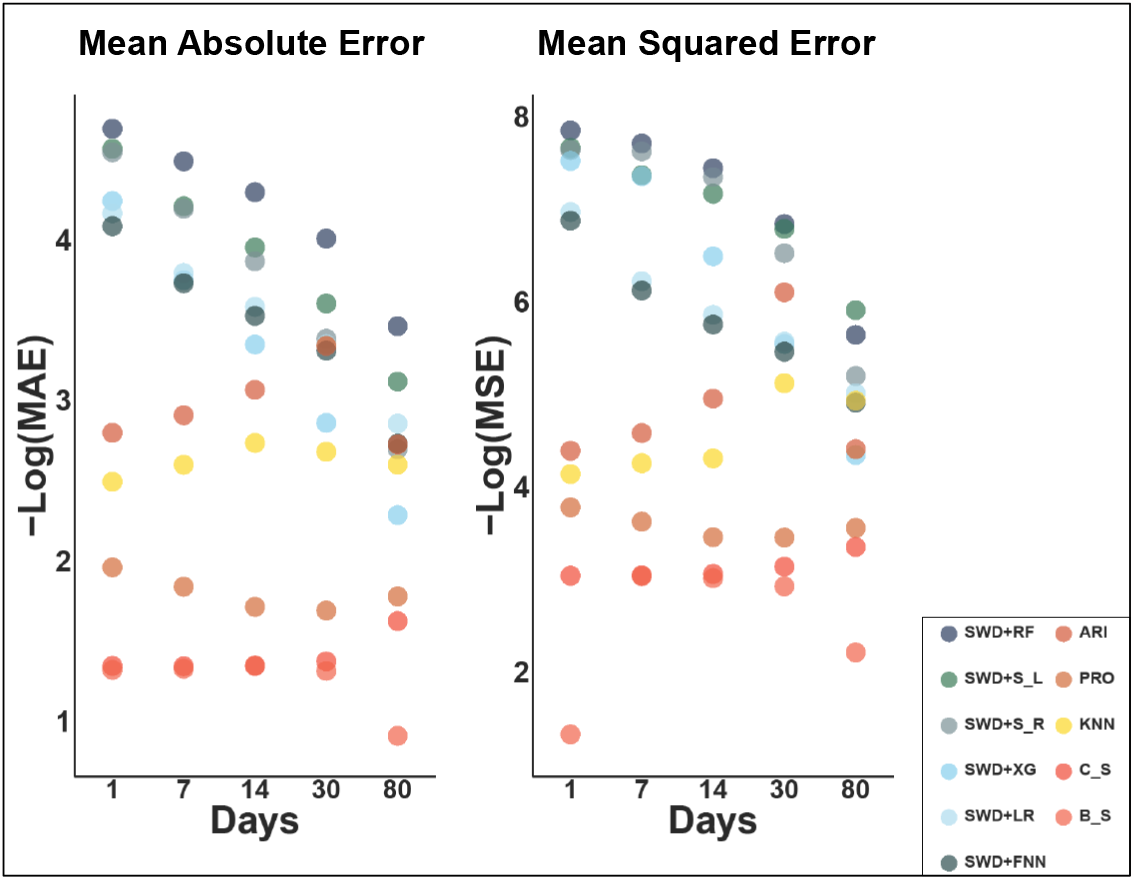
Visual comparisons of the performance of different learning methods in predicting the next 1, 7, 14, 30, and 80 days. On the left, the Mean Absolute Error (MAE) of these algorithms is displayed, and on the right, their Mean Squared Error (MSE). For clarity, we’ve applied the negative logarithm to both MAE and MSE scores. Supervised learning methods are represented in cold colors, contrasting with the warm colors used for time series models.

Next, we further compare the supervised learning and time series learning methods by taking a more refined look into the prediction accuracy. Figure 3 offers an in-depth look at each method’s performance. Similar to Figure 2, by training with mutation frequency data up to day *t*_0_ = 500, we predicted mutation frequencies for the subsequent 30 or 80 days and compared these predictions with actual data using scatter plots and barplots. In Figures 3A and 3C, scatter plots of observed versus predicted mutation frequencies reveal the extent of deviation for each method. Supervised learning methods are represented in cold colors, while time series methods are in warm colors. Among supervised methods, random forest (SWD+RF) and SVM (SWD+S_*L*_ and SWD+S_*R*_) demonstrate notable effectiveness, as evidenced by the overall placements of the points on the *x* = *y* axis. Interestingly, linear regression (SWD+LR) also shows commendable performance, surpassing time series methods, and provide comparable performance to XGBoost (SWD+XG), and forward neural network (SWD+FNN). Figures 3B and 3D quantify the deviation between predictions and observations for each of the 87 genomic locations from a discretized perspective. We assess how often predictions vary from actual values by 0.1%, 1%, and 5% over the next 30 and 80 days, using one of the best-performing method, SWD+RF. This visual analysis indicates that for most mutations, the prediction error for the upcoming 30 days is under 0.1%, and never exceed 1%. Even for the 80-day forecast, deviations exceeding 1% are rare and always within 5%. The results highlight the high reliability of 30-day predictions, especially with RF and SVM, as evidenced by scatter points closely aligning with the *x* = *y* axis. The majority of mutations show deviations below 0.1%, and all stay under 1%. For 80-day forecasts, despite higher deviations seen in the scatter plot (Figure 3C), linear kernel SVM emerges as the most accurate, and random forest also performed well, with almost all deviations remaining below 1% as shown in Figure 3D.

**Figure 3:**
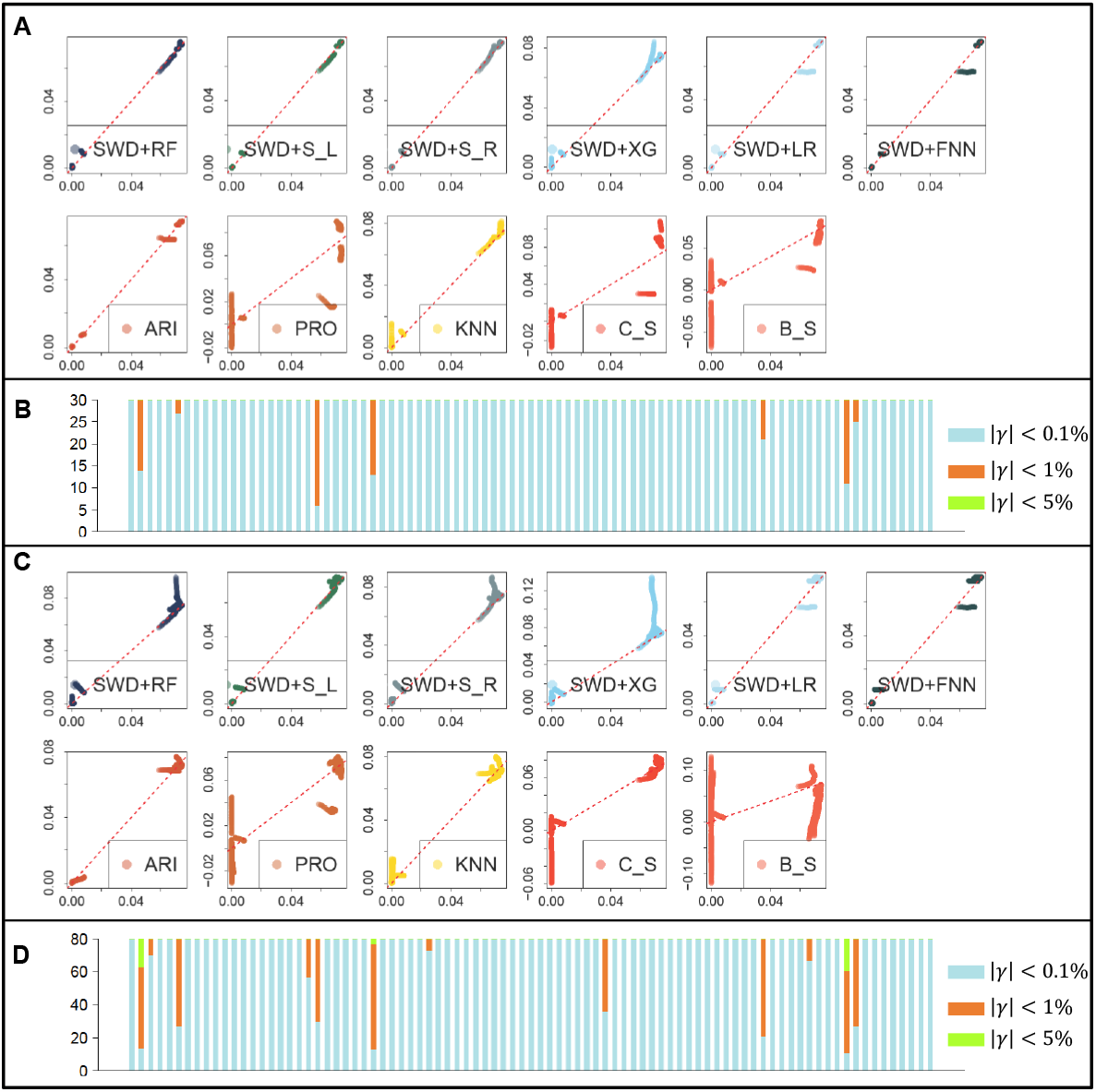
Comparisons of different learning methods in terms of prediction error for the future 30 and 80 days. A) Scatter plot of predicted (*x*-axis) and observed (*y*-axis) mutation rate for the next 30 days by gathering mutation frequency data of all 87 genomic locations. B) Barplot showcases the number of instances where the absolute difference between observed and predicted values is less than 0.1% (blue), 1% (orange), and 5% (green), respectively for predicting future 30 days using SWD+RF. C) Scatter plot for predicting future 80 days similar to A. D) Bar plot of level of deviations in predicting future 80 days similar to B.

The mutation frequency curves often exhibit numerous fluctuations, characterized by abrupt increases and declines, indicative of the unpredictable nature of viral mutation patterns (See Supplementary Figure S1). To assess the capability of the proposed methods in managing such irregularities, we selected two specific mutation curves, namely “`C_28311_T`” and “`AG_28878_TC`”. These two particular mutations are marked by a considerable number of segments exhibiting high fluctuations. For both mutations, we further handpicked seven segments that represent the fluctuations that spread throughout the whole time frame. We then evaluate how different learning approaches could handle the fluctuated mutation frequencies. For “`C_28311_T`”, shown in **Figure 4**, the bold curve represents its daily mutation rate, and the curve is mostly black, with seven red segments, and seven cyan segments each right after one red segment. For each supervised learning method, its model parameter is first trained using the all the mutation segments that arose from other mutation types excluding “`C_28311_T`”. To make prediction, the red segment, which is of length *d, d* = 21, serves as input predictors to predict the next day 1, day 2,…, until day 30 in a recursive fashion, i.e., the cyan segment; for time series methods, to predict each cyan segment, all the data points preceding this segment is used as training data to solve the model parameters, and prediction is directly made for the days within the cyan segment. Note the limitation with time series model which could not take other mutation types during training phase. The seven panel figures showcased the comparisons of predictions made by different learning methods with the true mutation rates, each for one handpicked segment, where the true mutation rate is marked by black dashed line. The supervised learning approaches (solid segments of cold colors) demonstrated much better predictions than the time series learning approaches (solid segments of warm colors), seen by their overall closeness to the observed segments (black dash segments). Specifically, when the segments are experiencing steep increase or decrease but with different rate (segments 5,6,7), time series learning approaches are often displayed to be less capable of catching up with the changing slope; When the segments are experiencing small to moderate changes, but are relatively fluctuating (segments 1,2,3), time series models tend to not able to capture the fluctuations. When the segments are relatively flat with very small rate of increase as compared with its preceding time segments (segment 4), most of methods perform comparatively, with some of supervised learning methods’ predictions lie closer to the observed truth in general. Among all the supervised learning methods, the two SVM methods have the overall best performance, by its ability to capture the fluctuations. We could draw similar conclusions for mutation “`AG_28878_TC`” in Supplementary Figure 2.

**Figure 4:**
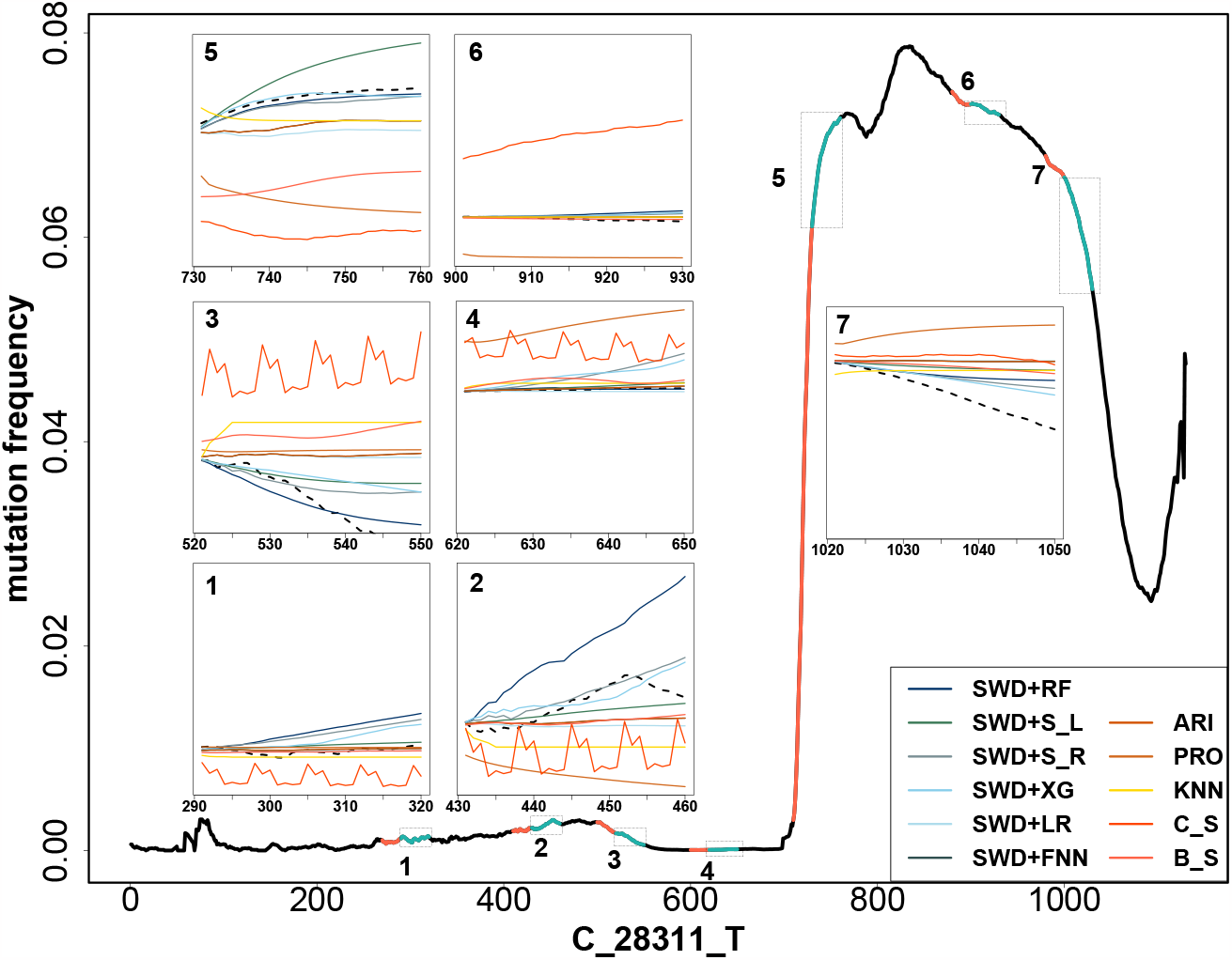
Prediction performance on seven future 30-day long segments of “G_28311_T”. Here on the mutation frequency sequence of “`G_28311_T`” (black line), we have selected 7 different temporal locations and 30-day long segments, each marked in blue on the whole mutation rate sequence plot. For each of the segment, we demonstrated the prediction performance of the 11 different algorithms on predicting future 30 days (seven panel figures).

In summary, while time-series models are effective in many forecasting situations, their ability to predict the evolving dynamics of mutations is limited. We identify three key reasons for this: 1) Time-series models excel at identifying consistent seasonal trends, which mutation patterns typically lack; 2) These models often train on each mutation independently, missing out on valuable inter-mutation information; 3) Time-series models generally face difficulties in making accurate long-term predictions in dynamic contexts such as mutation evolution. These limitations highlight the necessity for more sophisticated modeling techniques in complex forecasting scenarios, a point also emphasized by Coccia (2023) [28]. In contrast, our method demonstrates greater flexibility and applicability across different mutation types. It effectively captures the non-linear and intricate patterns hidden within mutation trajectories, offering more reliable predictions in these complex biological systems.

### 2.4 Performance evaluating in predicting unforeseen mutation patterns

In forecasting mutation frequencies, a pivotal task for any predictive model is its capacity to accurately anticipate future mutation frequency patterns, even for mutation types not encountered during training. To assess the adaptability of different modeling approaches to such scenarios, we divided the 87 mutation types into three predominant categories, and “simulated” scenarios in which a prediction method is trained on two categories of mutation types, subsequently applied to a distinct category for prediction. As the model has not been exposed to the mutation patterns from the category to be predicted, this simulates a scenario of forecasting unseen mutation types, thereby evaluating the model’s robustness and generalizability in predicting the evolution of novel mutations.

We have categorized the 87 genomic locations into three distinct groups based on their mutation patterns, as detailed in the Methods and Materials section. These groups are labeled as category 1, 2, and 3, comprising 63, 15, and 9 mutation types, respectively. Each category exhibits unique characteristics. Category 1 consists of mutation types where the highest mutation rate remains below 5%. Category 2 includes mutation types that peak earlier in the timeline, with a mutation frequency surpassing 5%. Category 3 features mutation types that exhibit their highest frequencies later in the timeline, with rates also above 5%. Similar to Figures 2 and 3, we demonstrate the methods’ performance in terms of MSE and MAE in Figure 5, and scatter plots and barplots in Figure 6.

**Figure 5:**
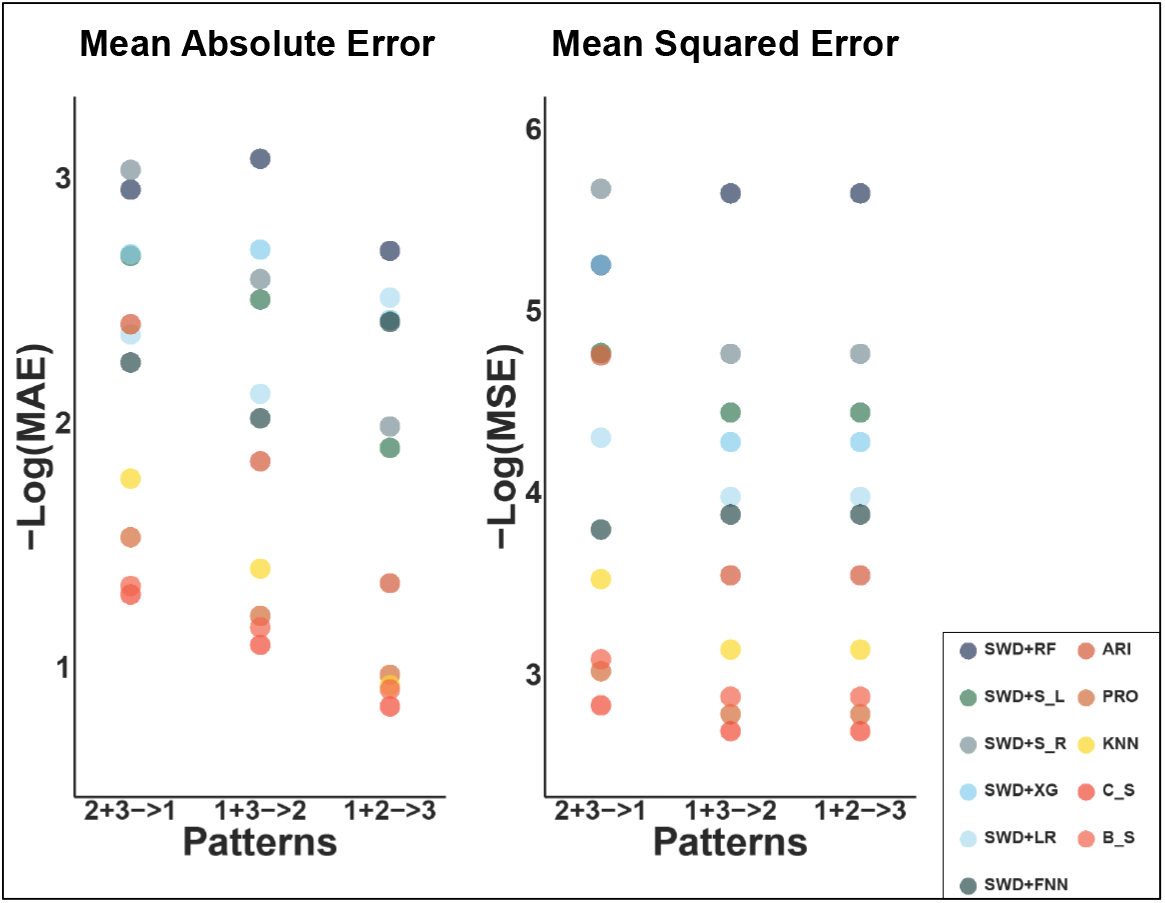
Robustness analysis. The prediction error measured by MAE (left) and MSE (right) are shown for predicting future 30 days under three different scenarios. For instance, (2+3→1) implies using pattern 2 and pattern 3 as the training set and pattern 1 as the test set. Similarly, cold and warm color bubbles indicate supervised and time series learning methods respectively.

**Figure 6:**
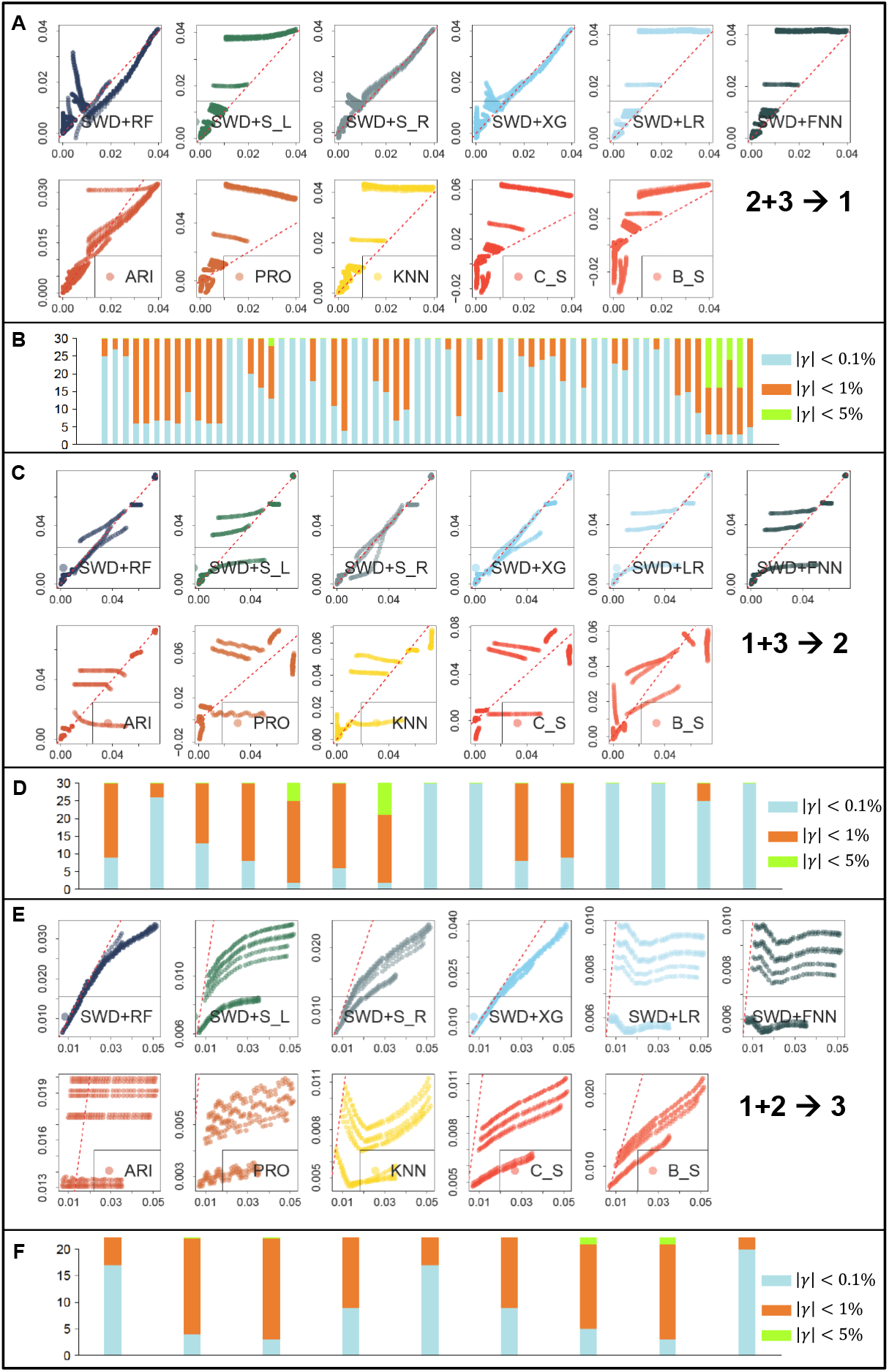
Robustness analysis. A) Scatter plot of predicted (*x*-axis) and observed (*y*-axis) mutation rate for the next 30 days when using pattern 1 and pattern 2 segments as the training data to predict mutation trend in pattern 3. B) Barplot of different levels of deviations between predicted and observed mutation rate for the next 30 days. C) and D) are similar to A) and B), but using pattern 1 and 3 to predict pattern 2, similarly for E) and F) which uses pattern 1 and 2 to predict pattern 3.

Specifically, for each supervised learning method, we trained each model using data from two of the three mutation categories and then tested its ability to predict the third. For instance, we used data from categories 2 and 3 to predict category 1, categories 1 and 3 for category 2, and categories 1 and 2 for category 3. Our prediction process was structured as follows: we selected a specific time point, *t*_0_, and aimed to predict mutation rates for the next 30 days (*t*_0_ + 1 to *t*_0_ + 30), using the preceding data (*t*_0_ *-d* + 1 to *t*_0_) as predictors. The training data, drawn from two categories, always preceded the prediction point *t*_0_. This method was applied consistently for predicting mutation frequencies in categories 1,2, and 3. We used random forest (SWD+RF) for generating the barplots. For our analysis, we set *t*_0_ = 500, a midpoint in our overall timeframe. In comparison, we also evaluated the performance of traditional time series models. Due to their limitation of modeling each sequence independently, the time series results were based on using data up to and including *t*_0_ to predict the mutation rates for the 30 days following *t*_0_ for each genomic location.

Figure 5 shows the performance of different methods in terms of MAE and MSE. It’s evident that for predicting unseen mutation types in categories 2 and 3, supervised learning methods significantly outshine all time series methods, as indicated by much lower MAE and MSE values. However, when forecasting for category 1, ARIMA (ARI) performs better than linear regression (LR) and forward neural network (FNN) in terms of both MAE and MSE, though it still lags behind the other four supervised methods. Here, predicting mutation patterns in category 1 poses a unique challenge, likely due to its larger number of mutation types and the generally flatter mutation rate curves compared to the other categories. These characteristics result in more distinct patterns, making accurate predictions more difficult.

Figure 6 offers an insightful view into the performance of these methods. Predicting unforeseen mutation patterns is inherently difficult, and this is reflected in the scatter plots (Figure 6 A, C, E). Here, almost all time series methods, along with some supervised learning methods like LR and FNN, show scattered, random predictions, with predicted and observed data points diverging significantly from the *x* = *y* axis. However, certain supervised methods, such as random forest (SWD+RF), SVM with a linear or radial kernel (SWD+S_*L*_ and SWD+S_*R*_), and XGBoost (SWD+XG), maintain more desirable performance, with most predicted and observed values closely aligning with the *x* = *y* axis. The barplots in Figure 6 (panels B, D, F) offer a more direct interpretation of each method’s absolute prediction accuracy. While there are more instances of higher prediction deviations (indicated by orange and blue colors) compared to the overall model trained on all mutation types (as seen in Figure 3), the majority of these deviations remain within 1%. Only in rare cases do the deviations exceed 1%, yet they still stay under 5%.

We have also analyzed the results when using raw frequency data instead of the first-order derivative data. Figure S3 in our study provides a clear illustration of the advantages of our data transformation approach compared to directly modeling raw frequency data. When using the raw frequency data, which neglects the inherent temporal order of the predictors, the results are significantly less precise. This is evident in the much more disorganized scatter plots shown in Figure S3 A, particularly when predicting mutation frequencies for the upcoming 30 and 80 days. The issue becomes even more pronounced when performing the robustness analysis which results in even greater disarray, lacking any clear order or pattern. Such findings emphasize the crucial role of our data transformation strategy and the use of first-order derivatives in preserving the sequential nature of the data, both of which are key factor in achieving accurate and reliable predictions.

Overall, these findings underscore the challenges in forecasting novel mutation patterns and highlight the superior performance of our supervised learning methods in managing these complexities, which is crucial for robust and reliable prediction models in virology and epidemiology.

## 3 Discussions and Conclusion

Our research on SARS-CoV-2 mutation patterns addresses a key element in understanding COVID-19’s progression. Influenced by viral mutations, immunity levels, social behaviors, and public health policies, the virus’s evolution presents intricate challenges. Predicting the realtime frequency and characteristics of these mutations is difficult due to their complexity and unpredictability. Traditional time-series models often struggle to capture the dynamic nature of these mutations, indicating a need for more refined approaches [29]. To address this need, we devised a supervised learning approach, employing a novel data transformation strategy, to forecast future mutations in real-time. Our results demonstrate the effectiveness of supervised learning methods, especially when modeling the first-order derivative of mutation frequencies.

The data transformation strategy we used is crucial. It allows the application of various advanced machine learning techniques, broadening the potential and adaptability of our model. By focusing on the first-order derivative rather than raw frequency, our method better preserves data sequence, crucial for accurately representing time series patterns. This machine learning approach has broader implications, suggesting potential for monitoring other infectious diseases and serving as a rapid-response tool in new disease outbreaks.

We developed comprehensive evaluation metrics, going beyond MAE and MSE to include visual and discretized measures of prediction deviation and testing performance under different fluctuation levels. Our robustness analysis was particularly crucial, simulating the challenge of predicting new mutation types not seen in training. This test is vital for evaluating the accuracy, flexibility, and generalizability of the models. Their ability to forecast the behavior of new mutations is essential for real-world applications, where predicting the virus’s evolution as new mutations emerge is critical.

Nevertheless, our methodology has limitations. Transforming time-series data for supervised learning can lead to a loss of temporal information, possibly obscuring important temporal patterns and relationships. Integrating time-based features or developing hybrid models combining time-series and supervised learning approaches could enhance our methodology. Additionally, the varying correlations among mutations add complexity. Advanced feature engineering and selection methods might be necessary to address these inter-mutational correlations effectively.

## 4 Methods and Materials

### 4.1 Data collection

We obtained all of our mutation rate data from the Global Initiative on Sharing All Influenza Data (GISAID) database. The GISAID database contains a large and rapidly growing collection of genomic data on the SARS-COV-2 from all over the world. We focused on the USA data only for several reasons: 1) The USA has experienced a diverse range of SARS-CoV-2 variants and has witnessed multiple waves of the pandemic, offering a rich and heterogeneous dataset that is reflective of the virus’s dynamic nature. 2) The USA has a well-established infrastructure for genomic sequencing and reporting, contributing to the high quality and reliability of the available data. The extensive geographical and demographic diversity within the USA provides a microcosm of varied environmental, societal, and healthcare contexts, enabling a comprehensive exploration of the factors influencing viral mutation. 3) The high incidence and prevalence of COVID-19 cases in the USA have resulted in a substantial volume of data, enhancing the statistical robustness of our analyses. Concentrating on this singular, yet diverse, dataset allows for a more controlled and nuanced investigation into the mutation patterns of SARS-CoV-2, while still capturing the complexity and variability inherent to the virus’s evolution.

The GISAID database encompasses not only the genomic sequences of the virus but also associated metadata, providing details such as the location and date of sample collection. Leveraging this resource, we compiled an exhaustive record of virus mutation events spanning from December 12, 2019 to January 26, 2023. In essence, each variant sequence was aligned to a reference genome, enabling the identification of the type of mutation at specific nucleotide positions, for instance, G*→*T, along with the precise genomic location of the nucleotide. Additionally, information including the geographic location (state and country) and the date of data collection for each patient’s virus, was also available. Each mutation event is designated based on the mutation type and its location; for example, ‘G 9053 T’ signifies a substitution of base G to T at location 9053 in alignment with the reference genome. The historical data of each mutation event form the basis for predicting its future mutation frequency. Note that despite the genome size of the virus being 30kb, most of the genomic locations have very low to zero mutation rate, and eventually, we have included 300 genomic locations that have mutation rate that peaked above 0.01. Among the 300 locations, many have mutation rate curve that are identical to others. After some filtering, we have come down to 87 locations with unique mutation rate curves. The method for calculating mutation rate is detailed below. The 87 locations were later categorized into three distinct groups based on two key criteria: the peak mutation rate and the timing of this peak. Curves that are always under 5% is category 1; curves that peak above 5% and before June, 2021 is category 2; and curves that peak above 5% and at or after June, 2021 is category 3.

### 4.2 Data preprocessing and transformation

To calculate the daily mutation frequency, we adopted a strategy akin to Mercatelli and Giorgi [30]. For any given day *i* and mutation *k*, we first identified the total count of samples collected during this time (*C*_*i*_), and the subset of these individuals who carry the mutation *k* (*c*_*ki*_). Thus, the daily mutation frequency was computed as *c*_*ki*_*/C*_*i*_. The raw daily mutations derived using this method exhibited a high degree of noise, hence we implemented a smoothing step to mitigate the impact of transient spikes and drops in the data. Basically, for any day *i*, the daily mutation is calculated as the average of day *i* and its next 9 days, i.e., average of a 10-day window. This smoothing step yields a more stable and representative depiction of mutation frequencies curve over time. We use the smoothed daily mutation frequency as *f*_*ki*_ throughout the article.

For time series approaches, they directly model the sequence of daily mutation frequency data for each individual mutation, with the objective being to learn a distinct curve for each and subsequently make predictions for future time frames. However, such approaches cannot integrate information across multiple mutation types and may struggle to generate accurate predictions for previously unseen mutations. Most critically, the inherent smoothing constraints of methods such as spline-based and autoregression techniques, including ARIMA and Prophet, may encounter difficulties when the time series does not exhibit a discernible seasonal trend that can be learned and extrapolated.

To overcome such limitation, we employed a novel data transformation approach such that we could employ the powerful machine learning methods. We assume that the mutation frequency follows certain Markov property, such that mutation frequency at day *i* depends on a window of *d* days that precede this date, i.e., *P* (*f*_*ki*_ = *f*_0_|*f*_*k,i-*1_, *f*_*k,i-*2_, …, *f*_*k*1_) = *P* (*f*_*ki*_ = *f*_0_|*f*_*k,i-*1_, …, *f*_*k,i-d*_). Utilizing this property, we devise an approach to cut the whole time frame into overlapping time segments each of length *d* + 1, and regard the last entry of the segment as response, and the preceding entries as predictors. Throughout this article, we treat *d* = 20. A large number of segments could be obtained such way, guaranteeing sufficient statistical power for the machine learning models. Training a machine learning model is find a function *g*^*\**^ such that *g*^*\**^ = argmin_*g*_∑_**s**∈**all** *𝒮*_ *d*_**s**_. Here *𝒮* denotes the set of training time segments, which could be obtained from all or part of the mutation types or time frame, and *d*_*s*_ evaluates the loss for one particular segment of length *d* + 1, i.e., **s** = (*s*_1_, …, *s*_*d*+1_), such as squared difference between predicted and observed rate for 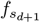.

Instead of looking for an optimal *g*^*\**^ for modeling the raw mutation rate, one could also look for an optimal *g*^*\**^ for modeling the first derivative of the mutation rate, which could better capture the inner dependencies of the predictors in 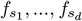 which possess a temporal order. Here, for any date *i* and mutation type *k*, the first order derivative, 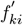, is approximated as the following: 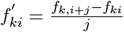. Here, we set *j* = 1. We have shown the superior performance of this strategy in Figure S3.

### 4.3 Constructing training and validation sets

To rigorously assess the performance of various methods, we partitioned the temporal mutation frequency data corresponding to 87 mutations into distinct training and testing datasets. This approach deviates from the traditional cross-validation technique, where training and testing instances are selected randomly. Instead, owing to the intrinsic temporal nature of our mutation frequency data, it is imperative to ensure that model training is confined exclusively to time frames prior to those for testing. This ensures that the model is not exposed to any mutation patterns from the testing data during training, maintaining the integrity and validity of our assessment. Specifically, for each of the 87 genomic locations each of 1,130 temporal observations, we obtained 1,130-*d* overlapping temporal segments. For example, segment 1 contains mutation frequency data from date 1 through *d* + 1, and segment 2 contains date 2 through *d* + 2, and so on. Note that here, the frequency data is the smoothed daily frequency data. We set *d* to be 20. The training and validation sets are constructed differently for different tasks. When measuring the overall performance (Figures 2, 3 and S3A), training data is obtained for all 87 genomic locations, but selecting only the time segments that are prior to date *t*_0_. Here *t*_0_ = 500. And the testing is performed for time horizon *t*_0_ + 1, …, *t*_0_ + *T*, for *T* = 1, 7, 14, 30, 80. When measuring performance regarding individual genomic location, namely “`G_28311_T`” and “`AG_28878_TC`” (Figure 4 and S2), training data is obtained for all other 86 mutation types excluding the one that is being tested. Here, the time segments is also confined to date prior to *t*_0_ = 500. Testing is performed for the seven hand-picked segments. When measuring performance regarding a large mutation type category (Figure 6A and S3B), training is obtained as time segments from two categories prior to *t*_0_ = 500, and testing is performed for the third category in regards to time horizon *t*_0_ + 1, …, *t*_0_ + *T*, for *T* = 30.

### 4.4 Parameter settings for various machine learning and time series methods

- Random Forest (RF): the R library `randomForest` was utilized. The key default parameters include a number of trees (`ntree`) set to 500, `mtry` determined as the square root of the number of input variables, `replace` set to TRUE, and `sampsize` set to the sample size.
- SVM with Linear Kernel (S_*L*_): The SVM with a Linear Kernel was implemented using the `e1071` library. The main default parameters include a misclassification cost (`cost`) of 1, `gamma` set to 1 divided by the data dimension, and `epsilon` set to 0.1.
- SVM with Radial Kernel (S_*R*_): The SVM with a radial Kernel was also implemented using the `e1071` library. The main default parameters are: misclassification cost (cost) set to 1, gamma set to 1 divided by the data dimension, and epsilon set to 0.1.
- XGBoost (XG): The XGBoost model was developed using the R library `xgboost`. The objective was set to “reg:squarederror”, the learning rate (eta) was set to 0.3, the maximum depth was set to 6, the evaluation metric was set to “rmse”, and the number of training rounds was set to 300. The key default parameters include the `booster` set to “gbtree”, L2 regularization weight (`lambda`) set to 1, and L1 regularization weight (`alpha`) set to 0.
- Linear Regression (LR): We used the builtin function `lm` in R for implementing linear regression.
- Feedforward Neural Network (FNN): The Feedforward Neural Network model was implemented using the `neuralnet` library. We set the hidden layers to 7 and each with 5 neurons respectively, and the linear output was set to TRUE. The key default parameters include an error threshold (`threshold`) of 0.01 and a maximum iteration step count (`stepmax`) of 1e+05.
- Autoregressive integrated moving average, ARIMA (ARI): The ARIMA model was automatically selected using the `auto.arima` function from the `forecast` package, with the input data being the length of the `train` dataset. By default, the function considers both non-seasonal and seasonal difference, with a maximum combined order of *p* + *q* + *d* of 5, and uses a stepwise selection approach.
- prophet (PRO): We implementd prophet using the `prophet` library. By default, it detects yearly and weekly seasonality, and automatically identifies potential trend change points.
- K-Nearest Neighbors (KNN): The forecasting was performed using the `knn forecasting` function. The model considered the previous 30 time points as lag variables, set by `lags = 1:30`. The number of nearest neighbors considered for the forecast was 50, as indicated by `k = 50`. The method used for forecasting was multi-input multi-output, specified by `msas = “MIMO”`.
- B-spline (B S): The B-spline basis functions were generated using the `bs` function from the `splines` package. The degrees of freedom for the B-splines were set to values in the vector `n0`, which includes 5, 10, 15, and 20. The linear model was then fitted using the `lm` function between the response variable and the B-spline coefficients.
- Cubic plines (C S): The `ns` function from the `splines` package was employed to generate natural spline basis functions. The degrees of freedom for the splines were specified by the vector `n0`, which includes the values 5, 10, 15, and 20. These degrees of freedom determine the complexity of the spline and the placement of knots in the data. The `lm` function was used to fit the model with `y` as the response variable and the spline coefficients as the explanatory variables.

## Supplementary Information

### Supplemental Figures

**Figure S1:**
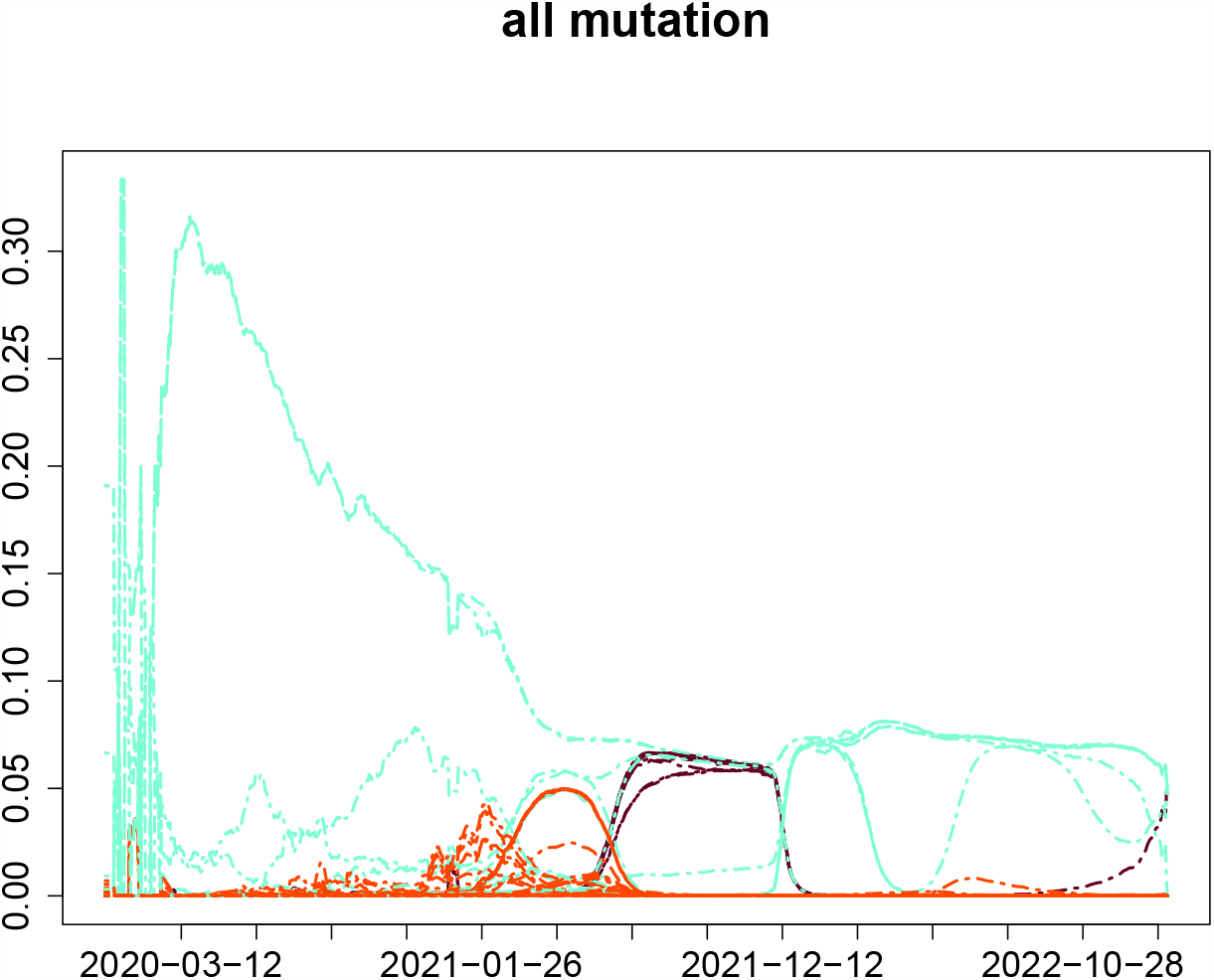
Mutation frequencies of all 87 mutation types, placed into three major categories, colored by cyan, orange red and maroon red. *x*-axis shows time, and the *y*-axis shows (smoothed) daily mutation frequencies.

**Figure S2:**
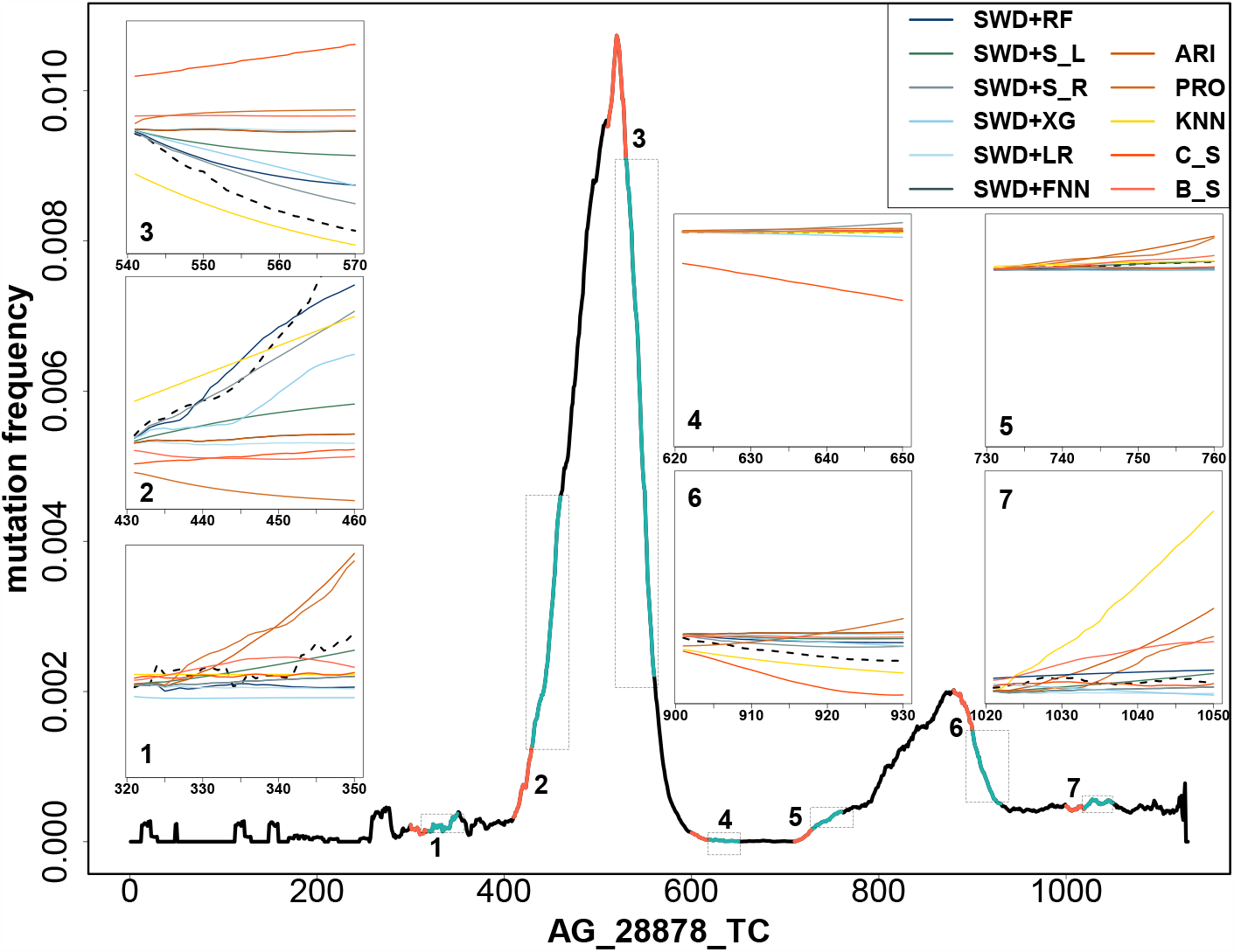
Prediction performance on several future 30-day long segments of “AG_28878_TC”. Here, on the mutation frequency sequence of “`AG_28878_TC`” (black line), we have selected 7 segments of 30-day length, each marked in blue on its whole mutation rate sequence plot. Similar to Figure 4, for each of the segment, we demonstrated the prediction performance of the 11 different algorithms on predicting future 30 days (seven panel figures).

**Figure S3:**
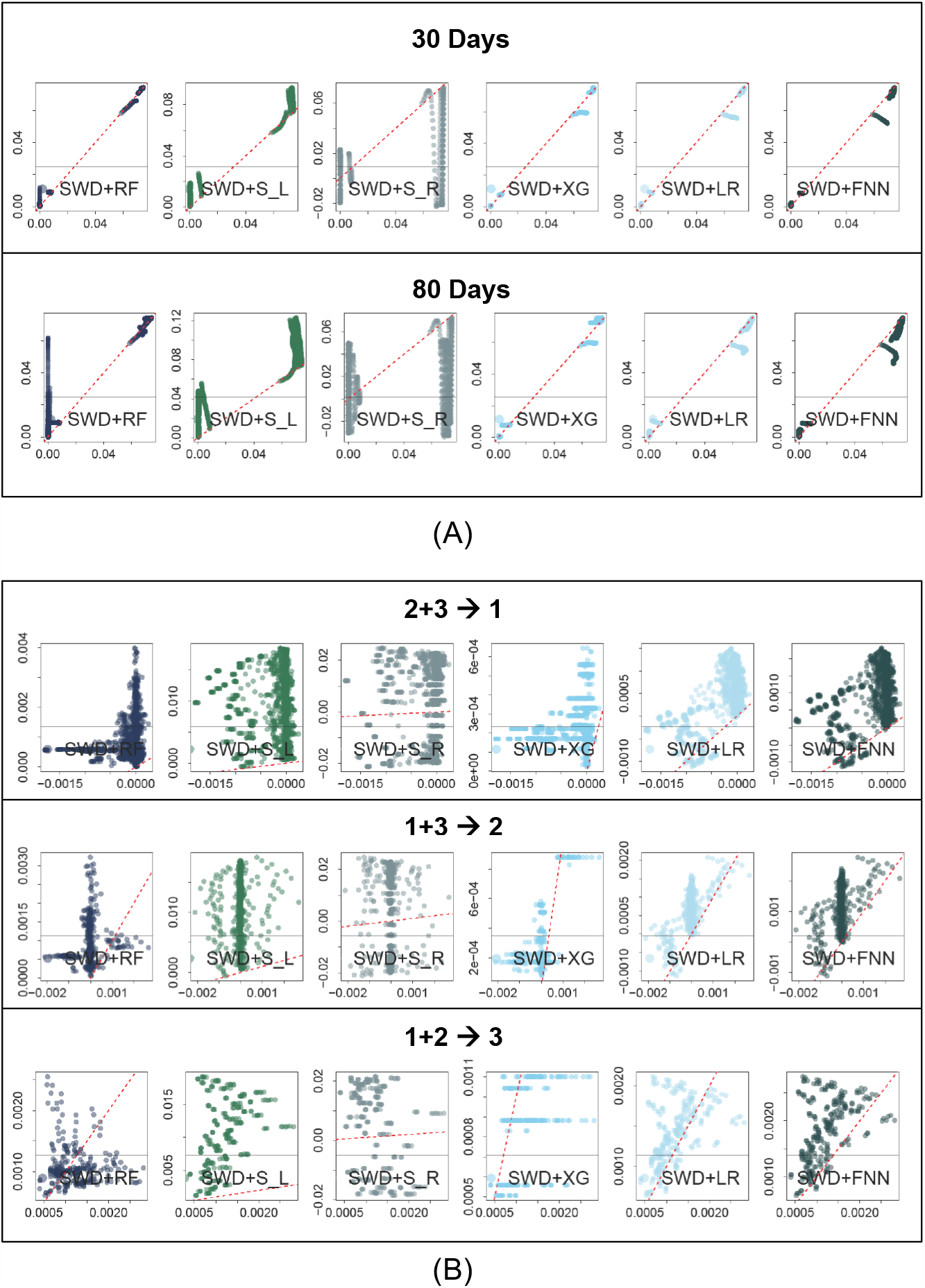
Prediction results using raw frequency data. (A) Scatter plot of observed (*y*-axis) and predicted (*x*-axis) mutation rate using raw frequency data for the next 30 (top) and 80 (bottom) days by gathering data of all mutations, similar to Figure 3A and 3C. (B) Robustness analysis for predicting the next 30 days by using two of the patterns as training and the rest as testing.

## References

[1] W. H. Organization. WHO Coronavirus (COVID-19) Dashboard. 2023. url: https://covid19.who.int.

[2] K. Tao, P. L. Tzou, J. Nouhin, R. K. Gupta, T. de Oliveira, S. L. Kosakovsky Pond, D. Fera, and R. W. Shafer. “The biological and clinical significance of emerging SARS-CoV-2 variants”. In: Nature Reviews Genetics 22.12 (2021), pp. 757–773.

[3] A. Telenti, E. B. Hodcroft, and D. L. Robertson. “The evolution and biology of SARS-CoV-2 variants”. In: Cold Spring Harbor perspectives in medicine 12.5 (2022), a041390.

[4] E. Hacisuleyman, C. Hale, Y. Saito, N. E. Blachere, M. Bergh, E. G. Conlon, D. J. Schaefer-Babajew, J. DaSilva, F. Muecksch, C. Gaebler, et al. “Vaccine breakthrough infections with SARS-CoV-2 variants”. In: New England Journal of Medicine 384.23 (2021), pp. 2212–2218.

[5] A. M. Carabelli, T. P. Peacock, L. G. Thorne, W. T. Harvey, J. Hughes, C.-1. G. U. C. de Silva Thushan I. 6, S. J. Peacock, W. S. Barclay, T. I. de Silva, G. J. Towers, et al. “SARS-CoV-2 variant biology: immune escape, transmission and fitness”. In: Nature Reviews Microbiology 21.3 (2023), pp. 162–177.

[6] W. T. Harvey, A. M. Carabelli, B. Jackson, R. K. Gupta, E. C. Thomson, E. M. Harrison, C. Ludden, R. Reeve, A. Rambaut, C.-1. G. U. (.-U. Consortium, et al. “SARS-CoV-2 variants, spike mutations and immune escape”. In: Nature Reviews Microbiology 19.7 (2021), pp. 409–424.

[7] R. P. Walensky, H. T. Walke, and A. S. Fauci. “SARS-CoV-2 variants of concern in the United States—challenges and opportunities”. In: Jama 325.11 (2021), pp. 1037–1038.

[8] Z. Chen, A. S. Azman, X. Chen, J. Zou, Y. Tian, R. Sun, X. Xu, Y. Wu, W. Lu, S. Ge, et al. “Global landscape of SARS-CoV-2 genomic surveillance and data sharing”. In: Nature genetics 54.4 (2022), pp. 499–507.

[9] M. Pachetti, B. Marini, F. Benedetti, F. Giudici, E. Mauro, P. Storici, C. Masciovecchio, S. Angeletti, M. Ciccozzi, R. C. Gallo, et al. “Emerging SARS-CoV-2 mutation hot spots include a novel RNA-dependent-RNA polymerase variant”. In: Journal of translational medicine 18 (2020), pp. 1–9.

[10] H. Lodish, A. Berk, S. L. Zipursky, P. Matsudaira, D. Baltimore, and J. Darnell. “Mutations: types and causes”. In: Molecular Cell Biology 4 (2000).

[11] K. Gangavarapu, A. A. Latif, J. L. Mullen, M. Alkuzweny, E. Hufbauer, G. Tsueng, E. Haag, M. Zeller, C. M. Aceves, K. Zaiets, et al. “Outbreak. info genomic reports: scalable and dynamic surveillance of SARS-CoV-2 variants and mutations”. In: Nature Methods 20.4 (2023), pp. 512–522.

[12] B. B. Oude Munnink, N. Worp, D. F. Nieuwenhuijse, R. S. Sikkema, B. Haagmans, R. A. Fouchier, and M. Koopmans. “The next phase of SARS-CoV-2 surveillance: real-time molecular epidemiology”. In: Nature medicine 27.9 (2021), pp. 1518–1524.

[13] J. Singer, R. Gifford, M. Cotten, and D. Robertson. “CoV-GLUE: a web application for tracking SARS-CoV-2 genomic variation”. In: (2020).

[14] A. T. Chen, K. Altschuler, S. H. Zhan, Y. A. Chan, and B. E. Deverman. “COVID-19 CG enables SARS-CoV-2 mutation and lineage tracking by locations and dates of interest”. In: Elife 10 (2021), e63409.

[15] J. Lu, L. du Plessis, Z. Liu, V. Hill, M. Kang, H. Lin, J. Sun, S. François, M. U. Kraemer, N. R. Faria, et al. “Genomic epidemiology of SARS-CoV-2 in Guangdong province, China”. In: Cell 181.5 (2020), pp. 997–1003.

[16] D. S. Candido, I. M. Claro, J. G. De Jesus, W. M. Souza, F. R. Moreira, S. Dellicour, T. A. Mellan, L. Du Plessis, R. H. Pereira, F. C. Sales, et al. “Evolution and epidemic spread of SARS-CoV-2 in Brazil”. In: Science 369.6508 (2020), pp. 1255–1260.

[17] L. Du Plessis, J. T. McCrone, A. E. Zarebski, V. Hill, C. Ruis, B. Gutierrez, J. Raghwani, J. Ashworth, R. Colquhoun, T. R. Connor, et al. “Establishment and lineage dynamics of the SARS-CoV-2 epidemic in the UK”. In: Science 371.6530 (2021), pp. 708–712.

[18] E. Volz, V. Hill, J. T. McCrone, A. Price, D. Jorgensen, Á. O’Toole, J. Southgate, R. Johnson, B. Jackson, F. F. Nascimento, et al. “Evaluating the effects of SARS-CoV-2 spike mutation D614G on transmissibility and pathogenicity”. In: Cell 184.1 (2021), pp. 64–75.

[19] F. Z. Najar, E. Linde, C. L. Murphy, V. A. Borin, H. Wang, S. Haider, and P. K. Agarwal. “Future COVID19 surges prediction based on SARS-CoV-2 mutations surveillance”. In: Elife 12 (2023), e82980.

[20] M. C. Maher, I. Bartha, S. Weaver, J. Di Iulio, E. Ferri, L. Soriaga, F. A. Lempp, B. L. Hie, B. Bryson, B. Berger, et al. “Predicting the mutational drivers of future SARS-CoV-2 variants of concern”. In: Science translational medicine 14.633 (2022), eabk3445.

[21] M. Khalid, D. Murphy, M. Shoai, J. N. George-William, and Y. Al-Ebini. “Geographical distribution of host’s specific SARS-CoV-2 mutations in the early phase of the COVID-19 pandemic”. In: Gene 851 (2023), p. 147020.

[22] S. Kim, Y. Liu, M. Ziarnik, S. Seo, Y. Cao, X. F. Zhang, and W. Im. “Binding of human ACE2 and RBD of omicron enhanced by unique interaction patterns among SARS-CoV-2 variants of concern”. In: Journal of computational chemistry 44.4 (2023), pp. 594–601.

[23] A. Ullah, K. M. Malik, A. K. J. Saudagar, M. B. Khan, M. H. A. Hasanat, A. AlTameem, M. AlKhathami, and M. Sajjad. “COVID-19 Genome Sequence Analysis for New Variant Prediction and Generation”. In: Mathematics 10.22 (2022), p. 4267.

[24] J. Zahradník, S. Marciano, M. Shemesh, E. Zoler, D. Harari, J. Chiaravalli, B. Meyer, Y. Rudich, C. Li, I. Marton, et al. “SARS-CoV-2 variant prediction and antiviral drug design are enabled by RBD in vitro evolution”. In: Nature microbiology 6.9 (2021), pp. 1188–1198.

[25] C. Bai, J. Wang, G. Chen, H. Zhang, K. An, P. Xu, Y. Du, R. D. Ye, A. Saha, A. Zhang, et al. “Predicting mutational effects on receptor binding of the spike protein of SARS-CoV-2 variants”. In: Journal of the American Chemical Society 143.42 (2021), pp. 17646–17654.

[26] M. Popovic. “Beyond COVID-19: Do biothermodynamic properties allow predicting the future evolution of SARS-CoV-2 variants?” In: Microbial risk analysis 22 (2022), p. 100232.

[27] J. L. Jacobs, G. Haidar, and J. W. Mellors. “COVID-19: challenges of viral variants”. In: Annual Review of Medicine 74 (2023), pp. 31–53.

[28] M. Coccia. “Sources, diffusion and prediction in COVID-19 pandemic: lessons learned to face next health emergency”. In: AIMS Public Health 10.1 (2023), p. 145.

[29] J. Chen, C. Gu, Z. Ruan, and M. Tang. “Competition of SARS-CoV-2 variants on the pandemic transmission dynamics”. In: Chaos, Solitons & Fractals 169 (2023), p. 113193.

[30] D. Mercatelli and F. M. Giorgi. “Geographic and genomic distribution of SARS-CoV-2 mutations”. In: Frontiers in microbiology 11 (2020), p. 1800.

